# Directed exploration in the Iowa Gambling Task: model-free and model-based analyses in a large dataset of young and old healthy participants

**DOI:** 10.1101/387019

**Authors:** Romain Ligneul

**Affiliations:** Donders Center for Cognitive Neuroimaging, Nijmegen, The Netherlands

## Abstract

The Iowa Gambling Task (IGT) is one of the most common paradigms used to assess decision-making and executive functioning in neurological and psychiatric disorders. Several reinforcement-learning (RL) models were recently proposed to refine the qualitative and quantitative inferences that can be made about these processes based on IGT data. Yet, these models do not account for the complex exploratory patterns which characterize participants’ behavior in the task. Using a dataset of more than 500 subjects, we demonstrate the existence of such patterns and we describe a new computational architecture (Explore-Exploit) disentangling exploitation, random exploration and directed exploration in this large population of participants. The EE architecture provided a better fit to the choice data on multiple metrics. Parameter recovery and simulation analyses confirmed the superiority of the EE scheme over alternative schemes. Furthermore, using the EE model, we were able to replicate the reduction in directed exploration across lifespan, as previously reported in other paradigms. Finally, we provide a user-friendly toolbox enabling researchers to easily fit computational models on the IGT data, hence promoting reanalysis of the numerous datasets acquired in various populations of patients.

## Introduction

Many disorders are associated with alterations of learning and decision-making. Therefore, standardized cognitive paradigms are increasingly used to improve diagnosis and treatments in these clinical populations. Developed 25 years ago (Bechara et al., 1994), the Iowa Gambling Task (IGT) remains one of the most popular tools used for this purpose (Fig. 1A). Over the years, it has been applied more or less successfully to many populations such as patients suffering from brain lesions, Parkinson disease, behavioral or substance addictions, mood disorders, personality disorders, etc. The IGT is thus of considerable importance for the development of scalable methods in cognitive science and in the emerging field of computational psychiatry.

**Figure 1.**
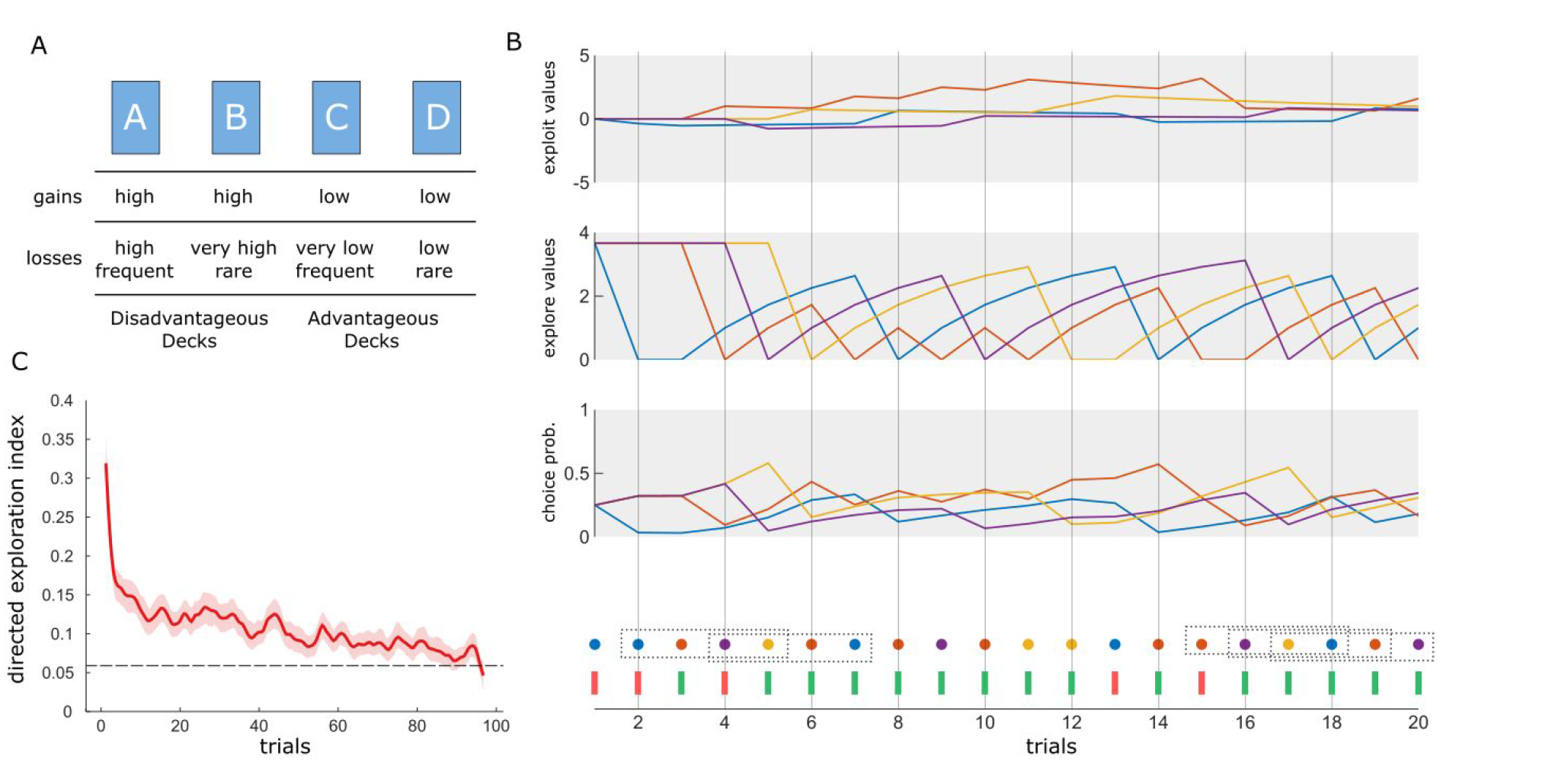
Directed exploration in the IOWA Gambling Task. (A) In the IGT, participants must sample 4 decks of card associated with gains and losses whose magnitude varies in a probabilistic manner. Unbeknownst to participants, decks C and D are advantageous despite offering smaller gains, because the losses are respectively very low or rarely encountered. (B) Here, we propose a new computational model accounting for trial-by-trial choices in the IGT. The “Explore-Exploit” (EE) model consist of two learning modules tracking respectively the net amount of money generated (exploit value, top) and the exploration value of each deck.

Yet, the classical analysis strategy for IGT data results in a crude estimate of decision-making deficits. Based on the relative preferences for “advantageous” decks (typically offering small gains but even smaller losses) over “disadvantageous” decks (typically offering big gains but even bigger losses), this approach does not leverage the full potential of the IGT. From a computational viewpoint, the IGT is indeed a highly complex task involving value-based learning, decision-making, working memory and different types of exploration. A series of computational models have thus been developed to better isolate these components, thereby offering clinicians and clinical neuroscientists more precise analytical tools to assess the cognitive profile of their patients. Computational neuroscientists interested in the IGT have first focused their efforts on the value-based learning and decision-making components of the task. Accordingly, the Expected Value (EV) or the Prospect Valence Learning (PVL, PVL-Delta) algorithms aimed at capturing distortions of reward and punishment processing. More recently, the Value Plus Perseveration (VPP) model was developed to capture systematic perseveration or alternation tendencies across successive decisions (Worthy et al., 2013). While the VPP model emerged as the most efficient model to describe choice data in the IGT, it can be criticized for its high number of free parameters (8) given the relatively short number of trials in the IGT (100) as well as for the uncertain cognitive validity of its perseveration module (Konstantinidis et al., 2014).

Here, we adopted another modeling strategy accounting for the existence of directed exploration (DE) in the class of multi-armed bandit tasks to which the IGT belongs (Collins and Koechlin, 2012; Daw et al., 2006; Raja Beharelle et al., 2015). Wilson and colleagues defined directed exploration as “a strategy in which choices are explicitly biased toward information”, as opposed to undirected (or random) exploration corresponding to a “strategy in which decision noise leads to exploration by chance” (Wilson et al., 2014). Thus, DE can refer to any regular choice pattern which: (i) maximize information about available options, (ii) cannot be readily explained by participants’ sensitivity to gains and losses. In the context of the IGT, a straightforward DE strategy is to allocate an “exploration bonus” to the behavioral options which are more uncertain, either because they have been sampled less often or because they have been sampled less recently than others.

Thus, we designed a compact computational architecture named Explore-Exploit in order to simultaneously capture exploitation, random exploration and directed exploration in the IGT using only 5 parameters. Besides random exploration which simply corresponds to decision noise (the opposite of decision consistency), the core innovation of this new model is thus to articulate two types of choice strategies: a reward-seeking strategy shaped by reinforcement history and an information-seeking strategy shaped by choice history (Fig. 1B). While the former is governed by the gains and losses delivered during the task, the latter depends on an exploration bonus specific to each participant, which can be either positive or negative depending on whether a given participant tends to explore or avoid options which have not been sampled recently.

In order to demonstrate the superiority of the EE model over existing alternatives, we reanalyzed an multi-study dataset of 504 participants who passed the standard version of the IGT (Steingroever et al., 2015a). State-of-the-art model comparison, simulation and parameter recovery analyses were performed to demonstrate the superior of the EE model. Second, in order to evaluate the cognitive validity to our model, we then focused on the data corresponding to the study by Wood and colleagues, which consisted in IQ-matched groups of old and young adults (Wood et al., 2005). Indeed, it was recently shown that directed exploration diminishes across lifespan (Mata et al., 2013; Schulz and Wu), so that the exploration bonus of older participants should be smaller than that of young participants. Third, we provide an open-source, user-friendly Matlab toolbox which has been developed to obtain the current results and which shall enable researchers who are not experts in computational models to re-analyze IGT data using both our new model and previous ones (see Appendix 1 and https://github.com/romainligneul/igt-toolbox).

These two first-order values are then summed and turned into a probability of choosing each deck. As such, the model implements a straightforward arbitration between reward- and information-seeking motivations. (C) The EE architecture was justified by the discovery of a peculiar choice pattern in the IGT. Namely, the probability of choosing 4 different decks within 4 consecutive trials (i.e a quantity referred hereafter as the directed exploration (DE) index) was largely above chance levels, especially during the beginning of the task.

## Results

### Presence of directed exploration in the IGT

First, we evaluated whether directed exploration (DE) occurred in the Iowa Gambling Task. To this end, we computed the DE index probing situations in which participants selected each of the four different decks over four successive trials using 25 independent consecutive quadruplets: e.g 1–4, 5–8, etc. In the 504 subjects dataset, we observed such pattern 1400 times (11.1%) while only 1182 occurrences would be expected under random exploration (i.e. 9.38%, binomial test: p<10^-10^). Note that this test is highly conservative, as the value-based exploration implemented by the softmax rule bias choices towards the most rewarded decks. Accordingly, a permutation approach in which trials were shuffled in time for each subject independently (total number of permutations: 5000) showed that the actual chance level was at 6.0%.

The target pattern was much more frequent in the first 20–30 trials of the task and it continuously declined as subjects formed more precise representation of each desk value and learned to exploit the reward structure of the task (Fig. 1C). Interestingly, DE had a complex but strong relationship with decision-making performances in the IGT. A GLM analysis indicated that subjects with the highest overall performance had lesser DE indexes (linear effect: t(1,501)=-3.40, p<0.001). However, we also observed low DE indexes in the worst subjects, which translated into a significant quadratic relationship between DE and performance (t(1,501)=2.13, p=0.034). Overall, the analysis of the DE index justified the development of the computational model capturing this important and previously overlooked exploration strategy in the IGT.

### Model comparison

The comparison of the EE architecture with the 4 aforementioned models was performed using the 504 subjects dataset. First, a fixed-effect analysis comparing summed Bayesian Information Criterion (BIC), Akaike Information Criterion (AIC) and Free Energy (F) metrics over the whole cohort demonstrated decisive evidence in favor of EE. In order to compare Free Energy with other metrics, it was transformed to -2*F for this analysis (Daunizeau et al., 2014). The difference between the EE model and other models was everywhere superior to 512 (Figure 2A; the least difference being observed with the VPP model based on the AIC metric). Note that a difference superior to 100 is generally considered as decisive evidence indicating that choosing the second-based fitting model would incur unacceptable information loss (Kass and Raftery, 1995). Going further, we performed a Bayesian Group Comparison (Stephan et al., 2009) based on the log-evidence of each model and treating model attribution as a random effect. In order to obtain log-evidences, we transformed AIC and BIC values to -AIC/2 and -BIC/2, respectively (Free Energy natively represents that quantity). Performed using all available metrics derived from the two fitting procedures (VB algorithm: BIC, AIC, F), this analysis showed that the estimated frequency of the EE Model was in every case superior to 50% and that its approximate exceedance probability (Ep) was always superior to 0.99 (Figure 2B). Overall, both approaches to model comparison provided overwhelming evidence in favor of the EE architecture, the only architecture able to account for directed exploration.

**Figure 2.**
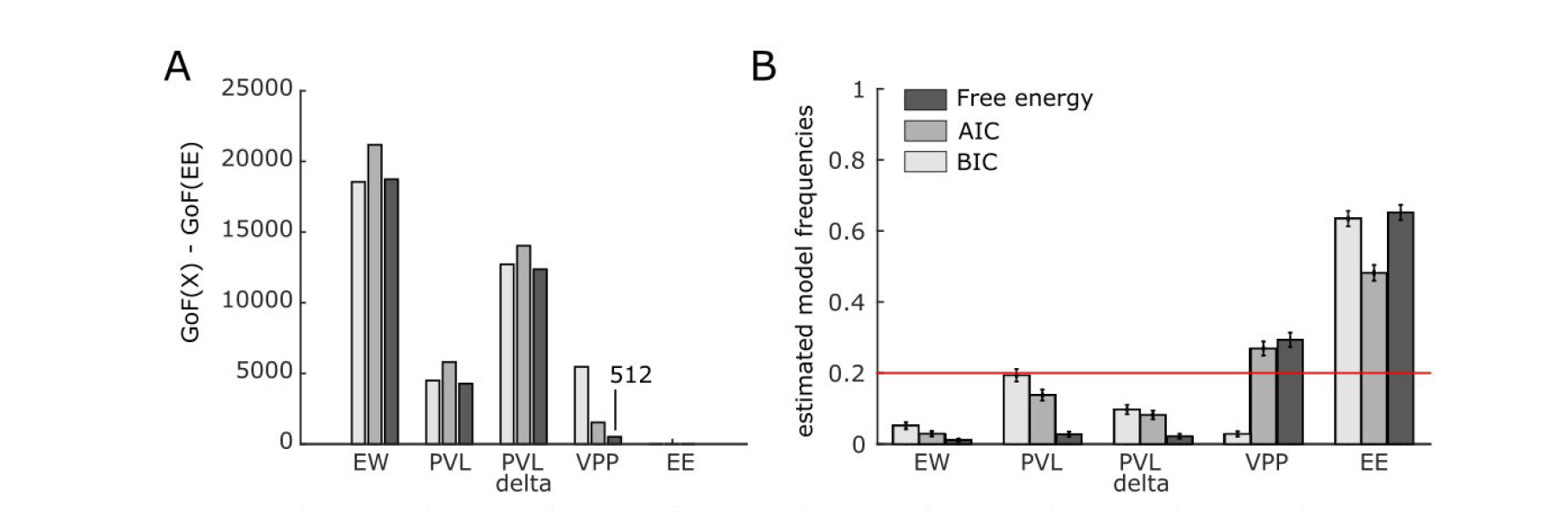
Model comparison. (A) Model comparison treating model attribution as a fixed effect showed that the EE model outperformed all other models on the 3 goodness of fit metrics. The least difference, observed with the VPP model using Free Energy, still reflected decisive evidence in favor of the EE model (Bayes Factor > 100)(Kass and Raftery, 1995). (B) Bayesian model comparison treating model attribution as a random effect also showed that the EE model outperformed all other models on the 3 goodness of fit metrics (exceedance probability superior to >0.99 in every case).

Regarding the relationship of model parameters with performance (defined as the number of advantageous minus disadvantageous deck selection), it appeared that value sensitivity was the strongest predictor (ρ=0.39, p<0.001), followed by *φ* (ρ=-0.24, p<0.001), decay (ρ=0.23, p<0.001) and temperature (ρ=-0.12, p=0.006)(Figure S1A). Moreover, although a substantial interindividual variability was observed in DE, the parameter *φ* corresponding to the exploration bonus was significantly superior to 0 (z=4.78, p<0.001). A correlation approach then confirmed that this key parameter reflected to which extent participants engaged in directed exploration (correlation DE index ~ *φ*: Spearman ρ=0.76, p<0.001). Since the likelihood of observing DE depends on which extent participants exploited the reward structure of the IGT, other parameters also predicted DE, but to a much lesser extent (update of exploration values: ρ=-0.26, p<0.001; value sensitivity: ρ=-0.20, p<0.001; inverse temperature: ρ=0.13, p=0.003)(Figure S1B).

### Simulation and parameter recovery

Next, we evaluated how the parameters estimated from individual choice data could be recovered for each model. Indeed, methodological studies in the field of computational modeling have demonstrated that different combinations of parameters can account for a given sequence of decisions, and that small deviations in parameters values can conversely result in significantly different sequences of decisions, hence impeding the interpretability of best-fitting parameters in some cases. Thus, we used the 504 sets of parameters associated with each model to simulate 504 *in silico* agents playing the IGT. For each deck, feedbacks were drawn randomly from their corresponding empirical distribution, hence keeping reward contingencies similar across actual and simulated tasks. Then, we applied the exact same fitting procedure to this simulated dataset. Parameter recovery was assessed by examining how the best-fitting parameters from this second-pass correlated with the best-fitting parameters from the first-pass (i.e that based the actual data).

Overall, the recoverability of parameters of EE was superior to that of other models (mean R=0.81, range: 0.67–0.95). EV and PVL also showed good recoverability (EV, mean=0.76, range: 0.66–0.83; PVL: mean=0.79, range 0.51–0.94), while PVL-delta and VPP were less stable (PVL-delta: 0.71, range: 0.5–0.86; VPP, mean=0.70, range: 0.41–0.94). In particular, it must be noted that the parameter *φ* reflecting the exploration bonus had the highest recoverability (0.95), hence making it a relevant target for the study of inter-individual differences (Figure 3A-E).

**Figure 3.**
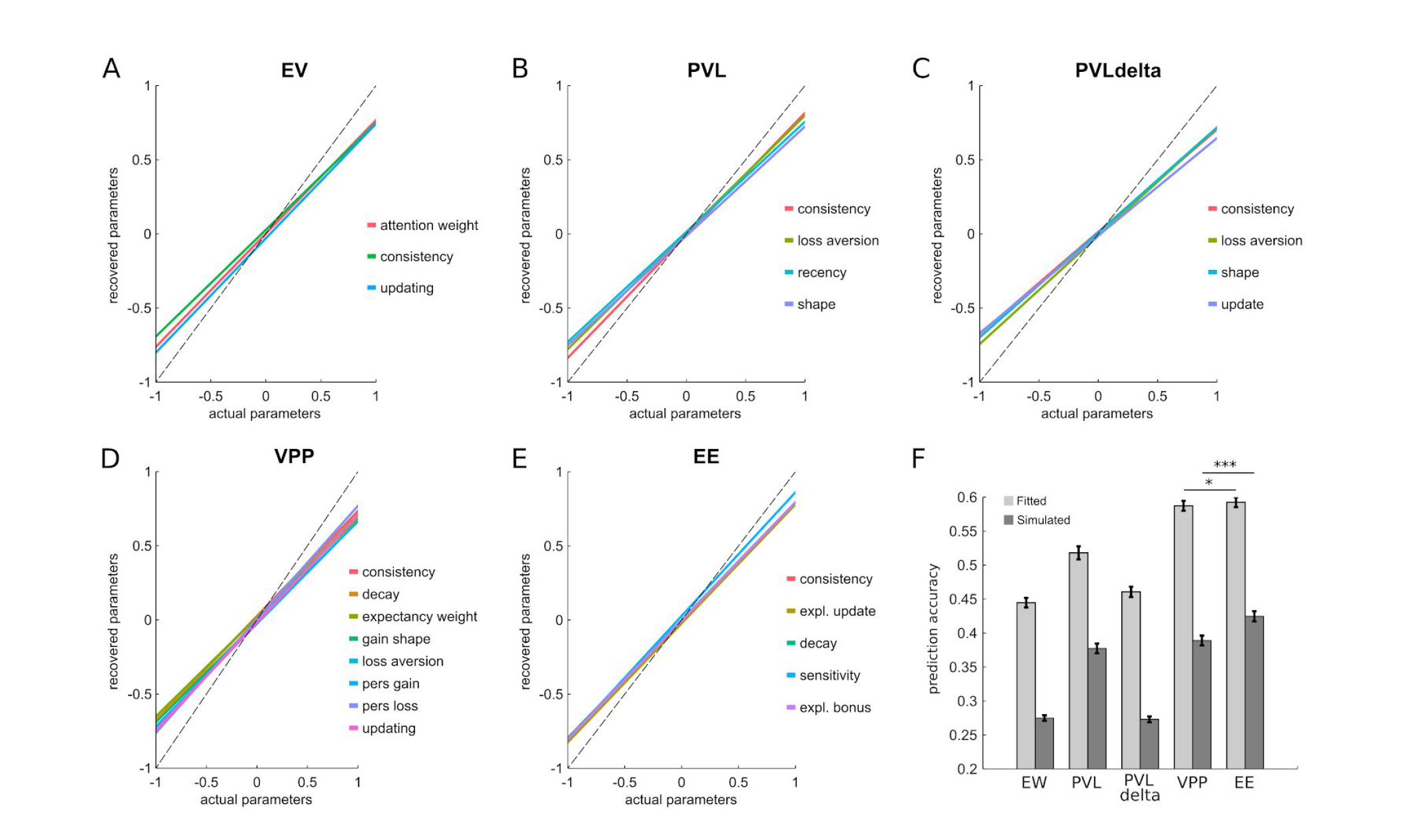
Simulation and parameters recovery. (A-E) The linear regression curves linking the parameters initially estimated based on participants’ choices and those estimated based on simulated choices showed that the EE model had the highest recovery performance (i.e minimal deviation from the identity diagonal). (F) The EE model was also the best model for predicting single decision, both for fitted and simulated choice data.

Then, we evaluated to which extent each model was able to reproduce participants’ choices using both first-pass and second-pass (i.e simulated) predictions (chance level: 25%, Fig. 3F). Again, the EE model was the most performant model. The first-pass (i.e fit on actual data) reproduced 59.2+/-16% of the choices, whereas the second best model in this respect (VPP) reproduced 58.7+/-17% of the choices. While the difference between EE and VPP was not significant (z=0.67, p=0.50), it must be noted that the VPP has 3 more free parameters which results in a greater chance of overfitting. In this respect, it is interesting to note the difference observed when comparing how the choices derived from simulated data reproduced participants’ choices. Here, the advantage of EE was clear, with 42.5+/-17% of successful predictions against 39+/-16% for VPP (z=3.63, p<0.001).

### Aging

In their study (included in the 504 participants dataset analyzed above), Woods and colleagues reported that old and young adults performed equally well on the Iowa Gambling Task, but resorted to different strategies. More precisely, old adults appeared to forget more rapidly about outcomes than healthy participants but compensated this forgetting by a better ability to translate what they learned into consistent choice patterns.

Thus, we used this subset of the data to evaluate how well the EE model could capture heterogeneities in IGT strategies and to validate our modeling approach (Figure 4A). In particular, based on the existing literature, we hypothesized that the exploration bonus should be lower in old as compared to young participants. First, we observed that old participants indeed forgot more rapidly than young participants according to the EE model, as indicated by a lower decay parameter (young: 0.55+/-0.27; old: 0.44+/-0.23; z=2.75, p=0.006). Second, the consistency parameter of old participants was indeed higher than that of young participants (young: 0.75+/-0.42; old: 0.92+/-0.41; z=2.71, p=0.007). Third and most importantly, young and old participants differed significantly in their *φ* parameter controlling the intensity of directed exploration in the IGT (young:0.94+:-2.14; old: 0.54+/-2.25; z=2.10, p=0.036). This latter result paralleled the model-free analysis of DE indexes which also revealed a reduction in directed exploration in the aging group (pattern frequency: young=16.5+/-14/6%, old=10.5+/-12.8%; t(151)=2.61, p=0.0099; Figure 4B) Overall, these results demonstrate the ability of the EE model to capture effects previously detected using simpler analysis strategies, as well the existence of age-related changes in directed exploration.

**Figure 4.**
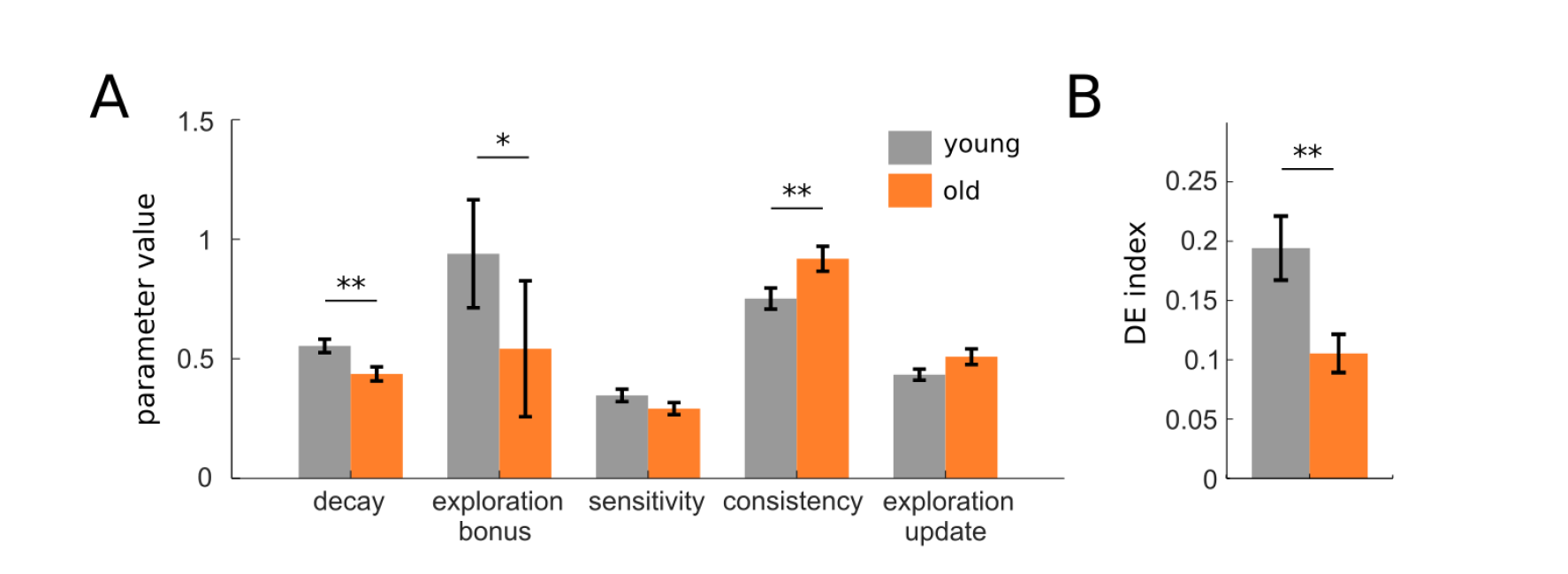
Model-based analysis of IGT behavior in a healthy aging cohort. (A) Using the EE model as a tool to refine the characterization of age-related changes in risky decision-making, we observed that old and young healthy adults differed on 3 parameters. Relative to young adults, old adults had a lower decay parameter (reflecting a faster forgetting of exploitation values), a higher consistency (reflecting a more deterministic choice policy) and a lower exploration bonus (reflecting a lower tendency to engage directed exploration). (B) The age-related reduction in the exploration bonus was reflected into a significantly lower DE index in old as compared to young participants.

## Discussion

In this study, we uncovered a new choice pattern reflecting the presence of directed exploration within the standard version of the IGT. Indeed, the selection of 4 different decks over 4 consecutive trials — a phenomenon captured by the “directed exploration” index — largely exceeded chance levels in a group composed of 504 participants, especially in the initial phase of the task. This discovery implies that the IGT might be used to study information-seeking behaviors within risky decision-making contexts. In order to better characterize and quantify this cognitive process within single individuals, we developed a new computational model (EE, for Explore-Exploit) able to articulate directed exploration with the motivation to optimize gains and losses, using only 5 parameters. The EE model outperformed the 4 most prevalent models previously used to fractionate the cognitive processes engaged by the IGT in terms of goodness of fit, prediction accuracy and parameter recovery. We further demonstrated the potential of this architecture to capture fine-grained differences in IGT behavior between young and old participants. Last but not least, we published the scripts used to generate our results under the form of a user-friendly Matlab toolbox which shall enable the community of researchers and clinicians relying on the IGT as a routine assessment of risky decision-making, in order to help them reporting more informative and detailed results with minimal programming and mathematical skills (see Appendix 1).

In the field of reinforcement-learning, most algorithms are oriented towards normative utility-maximization goals. To do so, they rely heavily upon reward prediction errors, a quantity widely used as a teaching signal enabling step-by-step convergence towards a utility maximum. Yet, likewise all gradient ascent methods, reinforcement-learning algorithms face the risk of reaching only local, rather than global, utility maxima. Directed exploration aims at solving this problem by expanding knowledge about the environment, despite immediate opportunity costs. Thus, directed exploration is particularly valuable when an agent is required to perform numerous decisions within complex or volatile contexts, in which the optimal policy may not be immediately obvious. The IGT is a canonical example of such environment. Indeed, with 4 possible actions leading to highly variable outcomes, the IGT is typically characterized by several successive phases: a “pre-punishment” period during which all decks only produce gains but no losses, a “pre-hunch” period during which punishments start occurring, a “hunch” period during which most healthy participants start feeling that the decks offering the highest average gains actually entails even higher average losses (i.e. deck A and B) and a “conceptual” period during which these participants are able to verbalize that the decks offering small gains (i.e. C and D) are actually the most advantageous ones (Bechara et al., 2005).

Although the term exploration is often applied to choices which are not maximizing utility according to a given computational model (a process now termed “undirected” or “random exploration”), the recent rise of predictive coding has recently completed this conceptualization (Clark, 2013; Friston et al., 2015; Huang and Rao, 2011). Indeed, this framework postulates that uncertainty-minimization constitutes a driving principle of our cognitive systems alongside utility-maximization, such that modeling the dynamics of exploratory decisions became an important endeavour in the field of reinforcement-learning. Accordingly, many recent studies have investigated uncertainty-minimizing behaviors in multi-armed bandit tasks using the Bayesian formalism. In this formalism, options whose mean value is the least precise (or, equivalently, associated with the largest variance) are the best candidates for exploration. Although the existence of uncertainty-driven exploration was confirmed by some of these studies (Badre et al., 2012; Boorman et al., 2009; Raja Beharelle et al., 2015), the type of directed exploration described here does not involve uncertainty computations. Instead, it relies on a simpler recency approach which promote the exploration of options which have not been selected for a while, independently of the objective uncertainty bound to their pay-offs. This methodology can be justified in three ways. First, despite its mathematical elegance, uncertainty-driven exploration does not provide a fully normative solution to the exploration-exploitation trade-off in multi-armed bandits (the process being heavily dependent upon higher-order priors regarding the structure of tasks). Second, the recency method implemented here might still reflect an uncertainty-based mechanism, if the subjective uncertainty associated with a given option increases with the duration elapsed since that option was tested for the last time. Third, the seminal study of Daw and colleagues had shown that uncertainty-driven exploration was not useful to describe exploratory patterns in a 4-armed bandit tasks sharing many commonalities with the IGT (Daw et al., 2006), hence keeping the door open for alternative algorithms.

The exploration module of our EE architecture helped going beyond existing models used to account for healthy participants’ decisions in the IGT. This differential performance contributes to ascertain the presence of directed exploration in the IGT. Combined with the model-free analysis of the DE index, this finding expand the heuristic value of the IGT beyond the study of exploitation and reward-seeking behaviors. Moreover, the fact that EE parameters were on average more recoverable than parameters of previous models facilitate the interpretation of inter-individual and inter-group differences. Numerous studies which had used the IGT to characterize clinical population may thus benefit from re-analyzing their data using the toolbox associated with the current paper. Once the trial-by-trial IGT data is converted to the appropriate Matlab format, this toolbox make such re-analysis extremely simple and intuitive, thanks to its compact but informative documentation and its densely commented scripts (see also Appendix 1). With minimal programming knowledge, the five models described hereinabove can be fitted to any standard IGT dataset, compared and evaluated with respect to parameter recoverability and prediction accuracy. These variables as well as other model-free measures (net scores, directed exploration indices, choice entropy, etc.) can also be calculated, plotted and compared across different groups.

Finally, in order to further validate our model and illustrate its utility for the analysis of group differences, we investigated how aging influenced EE parameters and more particularly the exploration bonus parameter. The results of this analysis were well aligned with those reported in the study of Wood and colleagues (Wood et al., 2005), in that the EE model still evidenced the exacerbated forgetting of previous outcomes in older adults, as well as the reduction in random exploration (i.e increased choice consistency) thought to compensate faster forgetting rates in these participants. More importantly, old adults also displayed a lower exploration bonus than young adults. This effect paralleled the reduction in directed exploration observed when computing directly the frequency of choosing 4 different decks over 4 consecutive trials (DE index). It is also highly consistent with recent papers showing that directed exploration reduces across lifespan (Mata et al., 2013; Schulz and Wu). Since directed exploration requires the retention of the last few choices made in the task, the phenomenon may be related to the decline of working memory performances sometimes observed in aging patients (Gazzaley et al., 2005).

At the neurobiological level, directed exploration likely depends on the prefrontal cortex (PFC), and more particularly on its rostrolateral portions (rlPFC). Indeed, several neuroimaging studies of directed exploration found that the rlPFC is more active during exploratory decisions (Badre et al., 2012; Boorman et al., 2009; Daw et al., 2006). Brain stimulation studies further showed that the disrupting or facilitating of rlPFC activity can significantly diminish or increase directed exploration, respectively (Raja Beharelle et al., 2015; Zajkowski et al., 2017). One may thus predict that disrupting rlPFC activities would similarly lower directed exploration and exploration bonuses in the IGT too. This involvement of the rlPFC might also explain the decrease in directed exploration seen in aging individuals, as grey matter density in this area is significantly reduced in old as compared to young adults (Tisserand et al., 2004). Yet, other neural systems certainly interact with the rlPFC to orchestrate information-seeking in reinforcement-learning tasks, including the dmPFC which may control the switch from exploitation to exploration strategies (Domenech and Koechlin, 2015/2). The prefrontal turn-over of dopamine might also play a pivotal role in regulating directed exploration (Frank et al., 2009; Kayser et al., 2015), whereas noradrenaline seems involved in the control of random but not directed exploration (Warren et al., 2017). Hopefully, investigating the wealth of data already existing for the IGT using the methodology proposed here might help elucidating more thoroughly the neural underpinnings of directed exploration in the near future.

To conclude, our study leveraged the power of an open “many labs” dataset in order to demonstrate the existence — and characterize the influence — of an overlooked behavior in the IGT. Building on previous work and more particularly on the Prospect Valence Learning (PVL) model (Ahn et al., 2008), the EE architecture represents not only a quantitative but also a qualitative improvement upon alternative models by shedding light on directed exploration. Besides enabling any experimenter to fit the EE and its ancestors (EV, PVL, PVL-Delta, VPP) on IGT data, the toolbox accompanying this paper might be used as an environment to develop even better models in the future. It must be acknowledged that this toolbox relies heavily on two other open-source packages for Matlab: modeling analyses largely depend on the VBA toolbox by Daunizeau and colleagues (Daunizeau et al., 2014) whereas visualizations take advantage on the Gramm toolbox by Morel (Morel, 2018). Last but not least, this study is fully aligned with the ideals of reproducibility and transparency in science: the dataset used is both large and freely available, while the scripts used to generate figures and statistics are available online alongside a clear documentation.

## Methods

### Dataset and participants

The dataset comes from a ‘many labs’ initiative grouping 10 studies and containing data from 617 healthy participants (Steingroever et al., 2015a). Here, we restricted the analysis to the subset of 7 studies which used the classical 100 trials version of the IGT, resulting in 504 participants (age range: 18–88 years old; for the 5 studies with available information about sex: 54% of females). Within this dataset, 153 participants come from a single study on aging (Wood et al., 2005). Among these participants, 63 are older adults (61–88 years old; 17 males) and 90 are younger adults (18–35 years old; 22 males) matched in terms of education level and intelligence (WASI vocabulary).

### Directed exploration index

In order to quantify directed exploration in the IGT, we computed the probability of choosing the 4 different decks during series of 4 consecutive trials. We refer to the frequency of such choice pattern as “DE index”. We used this metrics because the occurrence of such events has a probability of only 9.38% under purely random exploration (note that exploitation makes this probability even smaller by introducing an imbalance in the choice probability of different decks). Although directed exploration is certainty governed by more complex heuristics (resulting in more complex choice patterns), this index was used to ascertain its presence and provide an estimation of its intensity. Inferences about the presence of directed exploration used independent quadruplets of successive trials (i.e: 1–4, 4–8, etc.), whereas inferences about interindividual differences used dependent quadruplets to maximize sensitivity (i.e: 1–4, 2–5, 3–6, etc.).

### Previous models

Previous models have been exhaustively and excellently described in previous publications by Steingroever and colleagues (Steingroever et al., 2016). Therefore I will only provide a brief overview of their characteristics and then focus mainly on describing the features of the new EE model.

- The Expected Value (EV) model consists in a simple delta rule allowing for asymmetric consideration of gains and losses when updating the exploitation value of decks. It has thus a learning rate α (∈ [0,1]) parameter and a reward-punishment asymmetry parameter ω termed “attention weight” (∈ [0,1]).
- The Prospect Valence Learning (PVL) model is a working memory model in which past outcomes are discounted with a decay parameter δ (∈ [0,1]). Lower decay values imply faster discounting of past outcomes. Outcomes themselves are transformed based on the principles of prospect theory (Trepel et al., 2005): value sensitivity *v* and loss aversion ɭ (∈ [0,5]).
- The PVL-delta model applies the same transformation than the PVL model to outcomes, but it uses a delta rule to update exploitation values and therefore has no decay parameter but a learning rate (as in EV).
- Finally, the Value Plus Perseverance (VPP) model is the PVL-delta model extended with a “perserverance module”. This module has 3 parameter: a “persistance after gains” parameter ε_gain_ (∈ [0,1]), a “persistance after losses parameter” ε_loss_ (∈ [-1,1]), and decay parameter *δ*_pers_ (∈ [-1,1]) controling to which extend past persistance values are discounted. At the decision stage, persistance values and exploitation values are combined thanks to an expectancy weight parameter ω_Ev_ (∈ [0,1]).

All the models described above have in a addition a consistency parameter determining to which extend choices are driven by learned values (or any type). This consistency parameter *c* is allowed to fluctuate in the [0,5] interval and is transformed before being used as an inverse temperature parameter β (β = 3*^c^*–1), except for the EV model where c is allowed to fluctuate in the [-2,2] interval and is transformed differently (β = (t/10)*^c^* with t corresponding to current trial number). In sum, the EV model has 3 parameters, the PVL and PVL-delta models have 4 parameters and the VPP model has 8 parameters. Note that the consistency parameter capture the exact opposite of “random exploration” which corresponds to decision noise (or temperature).

### Explore-Exploit model

Like all other models, the Explore-Exploit (EE) model updates “exploitation values”, keeping track of the recent trends in gains and losses associated with each deck. Like the VPP model, the EE model also consists of two separate modules learning different values for each deck. However, the EE model differs from the VPP model in that it does not update “perseveration values” after gain and losses but a single “exploration value” for each deck, which depends only upon choice (rather than gains/losses) history. On each trial, these two values are simply summed in a deck-wise fashion before being transformed by a conventional softmax step into choice probabilities. Hereafter, we describe the seven equations and the five parameters which fully characterize EE.

The exploitation module is inspired by the PVL model (Steingroever et al., 2013) although it includes no “loss aversion” parameter: value sensitivity, controlled by θ (bound between 0 and 1), is instead applied to both wins and losses.

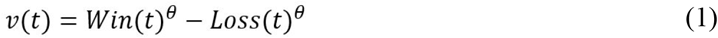

On each trial, the exploitation value of each desk d is updated according to the following equations:

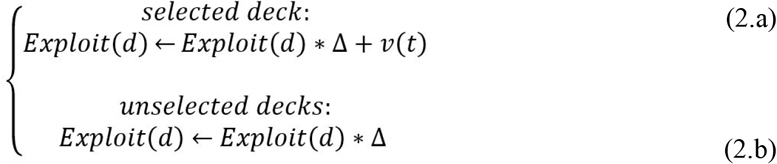

Equation (2.a) controls the update of the deck chosen, by adding the feedback just experienced to the (decayed) value of this deck. Equation (2.b) controls the update of unchosen decks, whose exploitation value progressively returns to 0 at a rate controlled by the decay parameter Δ (bound between 0 and 1). Note that a decay of 1 indicate that exploitation values are integrated over all previous trials, while a decay parameter of 0 indicate that subjects decisions rely mostly on the last feedback obtained in each deck.

The main innovation provided by EE consists in modeling directed exploration in the IGT. Exploration values reflect the attractiveness of each deck as a function of the number of trials for which the deck has *not* been selected, implying that exploration values are agnostic regarding the monetary feedbacks experienced in the task. As such, they capture a pure information-seeking process, hence contrasting with Bayes-based uncertainty-minimization algorithms as well as random exploration modeled by the softmax temperature or e-greedy rules (Daw et al., 2006). Exploration values are thus controlled by the following equations:

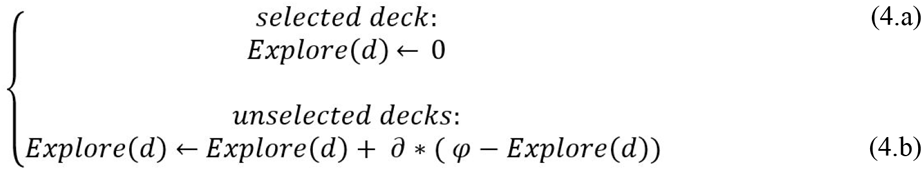

Equation (4.a) controls the update of exploration values for the selected deck, which fall to zero as soon as the outcome of that deck is sampled. Equation (4.b) controls the update of unselected decks, which is governed by a simple delta-rule. The learning rate γ (boundy22 between0and 1) determines at which speed the exploration values return to the initial value of the exploration bonus *φ* (unbounded). A positive exploration bonus implies that the participant is attracted by decks which have not been explored recently, whereas a negative exploration bonus implies that the participant tends to favor familiar decks. Importantly, the exploration bonus *φ* therefore reflects the strength of directed exploration, so that a more positive value will translate into higher probability of reproducing the aforementioned pattern of 4 different choices over 4 consecutive trials (all being equal otherwise).

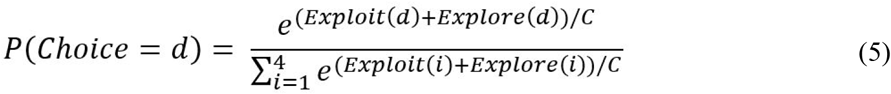

Finally, equation (5) models decision-making as a stochastic process influenced by the consistency parameter C: a higher C value indicates that choices are strongly driven by the composite values derived from Equations 1–4, whereas a C value of zero indicate random selection of each deck. Note that C results from the transformation of an inverse temperature β (bound between 0 and 5), in order to match PVL, PVL-Delta and VPP models (where C = 3^β^ -1 as well).

The architecture of EXPLORE can account for purely random exploration (β=0), for purely value-based exploitation (β>>0, θ>0 and *φ*=0), for purely directed exploration (β>>0, θ=0, and *φ*>0) and for a mixture of value-based exploitation, directed exploration and random exploration. Note that under purely directed exploration (β>>0, θ=0, and *φ*>0), the model predicts that the 4 decks should be successively selected in a cyclical manner during the whole task (e.g. 3,2,4,1,3,2,4,1,3,etc.). Indeed, on each trial, the deck with the highest exploration value would always be the deck which has not been selected for the longest period of time.

### Fitting procedures

A validated toolbox (http://mbb-team.github.io/VBA-toolbox) was used to optimize model parameters (Daunizeau et al., 2014). This toolbox relies on a Variational Bayesian (VB) scheme. Compared to non Bayesian methods, this approach has the advantage of accounting for the uncertainty related to estimated model parameters and of informing the optimization algorithm about prior distributions of parameters’ values. All priors were innately defined as Gaussian distribution of mean 0 and variance 3, which approximates the uniform distribution over the [0–1] interval after sigmoid transformation. Depending on the range of values in which each parameter was allowed to vary, the sigmoid-transformed parameters were further stretched or shifted to cover different intervals while preserving the flatness of their prior distribution (e.g. “multiplied by 2, minus 1”, to obtain the interval [-1,1]). Model comparison results were replicated using a non-Bayesian model fitting procedure which relied on the standard *fminunc* function of Matlab (line-search algorithm).

All hidden states (i.e values) were initialized at 0 except for exploration values which were initialized at 1 (since no deck has been sampled at the beginning of the task). The VB algorithm was not allowed to update the initial values for hidden states.

### Model comparison

Comparison of EE model with the 4 alternatives was first based on a classical fixed-effect analysis comparing summed Bayesian Information Criterion (BIC), Akaike Information Criterion (AIC) and Free energy (F) metrics over the whole group. In this approach, it is classically considered that a difference of 10 units between the models with the lowest and the second lowest criterion value reflects very strong evidence in favor of the model with lowest value (corresponding to a Bayes Factor of 150).

Then, a Bayesian Group Comparison was performed which treated model attribution as a random-effect varying from subject to subject. Also based on BIC, AIC and F, this type of analysis produces an exceedance probability corresponding to the probability that a given model is more likely than any other candidate model (Stephan et al., 2009).

### Parameter recovery and simulation

There is a growing consensus among computational neuroscientists that evaluating models only based on goodness of fit metrics like the AIC or BIC is not sufficient (Palminteri et al., 2017; Steingroever et al., 2015b). The problem is particularly salient when one aims at drawing inferences about cognitive processes from estimated parameters (which is most often the case), because the same choice pattern can sometimes be explained by very different combinations of parameters and because models with better goodness of fit metrics do not always better reproduce qualitative choice patterns. To address these issues and ensure that the EE model performed equivalently or better than the VPP model in this respect, we performed simulation and parameter recovery analyses detailed below.

We used the best-fitting parameters of each subject to simulate an artificial decision-maker confronted to the IGT. Simulated choices were generated stochastically according to the consistency parameter, and feedbacks (gains/losses) were drawn from the distributions of feedbacks actually encountered by the participants. Then, we reran model estimations based on these simulated choicesn, which resulted in a new set of parameters. The quality of parameter recovery for the EE and VPP models could then be assessed by examining the correlation of this second set of parameters with the parameters initially obtained by fitting real choices. Finally, we examined to which extent the initial choices predicted by the model and the choices performed by the simulated participants matched the actual choices of the participants, across models. In this latter analysis, we restricted our statistical inference and compare the EE model with the second-best fitting model only.

## Acknowledgements

RL would like to thank Roshan Cools, Guillaume Sescousse and Zachary Mainen for their support, as well as the Fyssen Foundation who funded RL during the preparation of this manuscript.

